# Isolation and characterization of a neoxanthin synthase gene functioning in fucoxanthin biosynthesis of *Phaeodactyum tricornutum*

**DOI:** 10.1101/2022.10.18.512692

**Authors:** J.K. Sui, Z.Y. Zhang, J.C. Han, G.P. Yang, T.Z. Liu, H. Wang

## Abstract

Golden-brown xanthophyll fucoxanthin in marine organisms, especially in diatoms, has attracted widespread attention because of its diverse biological activities. However, the biosynthetic pathway of fucoxanthin remains unclear in diatoms. Fucoxanthin may derive from either neoxanthin or diadinoxanthin pathway. However, the key point is whether neoxanthin and its synthesizing genes exist or not. In this study, we successfully identified a few xanthophylls in trace amounts in the concentrated fraction of carotenoids of diatom *Phaeodactylum tricornutum* cultured at different light intensities with the co-chromatography method, and cloned the neoxanthin synthase (NXS) gene which was not annotated in diatom genome. The *NXS* knockdown and knockout experiment show a positive correlation in the accumulation of neoxanthin and zeaxanthin while a negative correlation in violaxanthin and fucoxanthin with the expression of *NXS. In vitro* assay evidenced that neoxanthin is the precursor for fucoxanthin biosynthesis, indicating that other molecules intermediate the conversion between violaxanthin and fucoxanthin. Overall, we cloned a novel gene functioning in neoxanthin biosynthesis, which should aid to clarifying the fucoxanthin biosynthetic pathway in diatom.

## 1. Introduction

Microalgae are photosynthetic organisms that utilize carbon dioxide to synthesize high-value phytochemicals, making economic, environment-friendly and sustainable manufacturing of these chemicals possible. Diatoms are among the most successful microalgal cell factories and contribute about 40% of marine primary productivity. They are also commercially cultured to produce fucoxanthin, a specific non-provitamin A carotenoid (Butler et al., 2020). Fucoxanthin is also the partner of the fucoxanthin chlorophyll a/c-binding proteins (FCPs) of the unique light-harvesting antenna with exceptional light-harvesting and photoprotecting capabilities. It acts as a functional xanthophyll, transferring excitation energy to chlorophyll of the photosynthetic reaction centers (Domingues et al., 2012; Papagiannakis et al., 2005). Fucoxanthin has a unique molecular structure with an allenic bond, a conjugated carbonyl, a 5,6-monoepoxide and an acetyl group, which endows fucoxanthin with a variety of nutraceutical bioactivities and anti-oxidation, anti-cancer, anti-diabetic, anti-inflammatory and anti-metastatic effects (Bertram, 1993; Gómez-Loredo et al., 2016; Peng et al., 2011; Satomi, 2012). Diatoms grow fast and have been considered as the potential industrial producers of fucoxanthin. Unfortunately, the biosynthetic pathway of fucoxanthin in diatoms remains unclear.

Currently, fucoxanthin biosynthesis in diatoms has been explored physiologically, which may derive from geranylgeranyl pyrophosphate (GGPP). Several genes involved in its synthesis have been identified in comparative genomic analysis (Bowler et al., 2008). The condensation of two GGPP molecules catalyzed by phytoene synthase (PSY) forms the phytoene as a 15-cis isomer, which is then converted into lycopene via a serial desaturation and isomerization. Lycopene is cyclized by ε-LCY or β-LCY into α-carotene or β-carotene. The primary steps from GGPP to β-carotene are common among all species. Beta-carotenoid is first hydroxylated to form zeaxanthin by β-cryptoxanthin and β-hydroxylase (β-OHase) (Tian and Dellapenna, 2001), and then zeaxanthin epoxidase (ZEP) catalyzes zeaxanthin in two consecutive steps to yield antheraxanthin and violaxanthin (Jahns and Holzwarth, 2012). However, the final step of fucoxanthin biosynthetic pathway that start with violaxanthin remains unclear in diatoms.

Several hypothetical pathways have been proposed and discussed (Fig. 1) (Kolber et al., 1988). The neoxanthin pathway may involve the conversion of vioxanthin to neoxanthin, which is the branching precursor for the formation of both diadinoxanthin and fucoxanthin. Neoxanthin presents in the light-harvesting complexes (LHCs) of high plants and algae (Mikami and Hosokawa, 2013), constituting the violaxanthin cycle as a balancer responding to a wide range of light intensities in association with zeaxanthin and violaxanthin (Havaux and Niyogi, 1999). There is a close biosynthesis link between the allenic double bonds and acetylenic bond from neoxanthin to diadinoxanthin (Dambek et al., 2012). Violaxanthin may also be converted into diadinoxanthin, a precursor of fucoxanthin, because the diadinoxanthin cycle is an important bioprocess involved in the biosynthesis of light-harvesting xanthophylls (Lohr and Wilhelm, 1999). Therefore, the identification of neoxanthin and relationship verification between neoxanthin and fucoxanthin are crucial for fucoxanthin biosynthesis pathway elucidation.

**Fig. 1.**
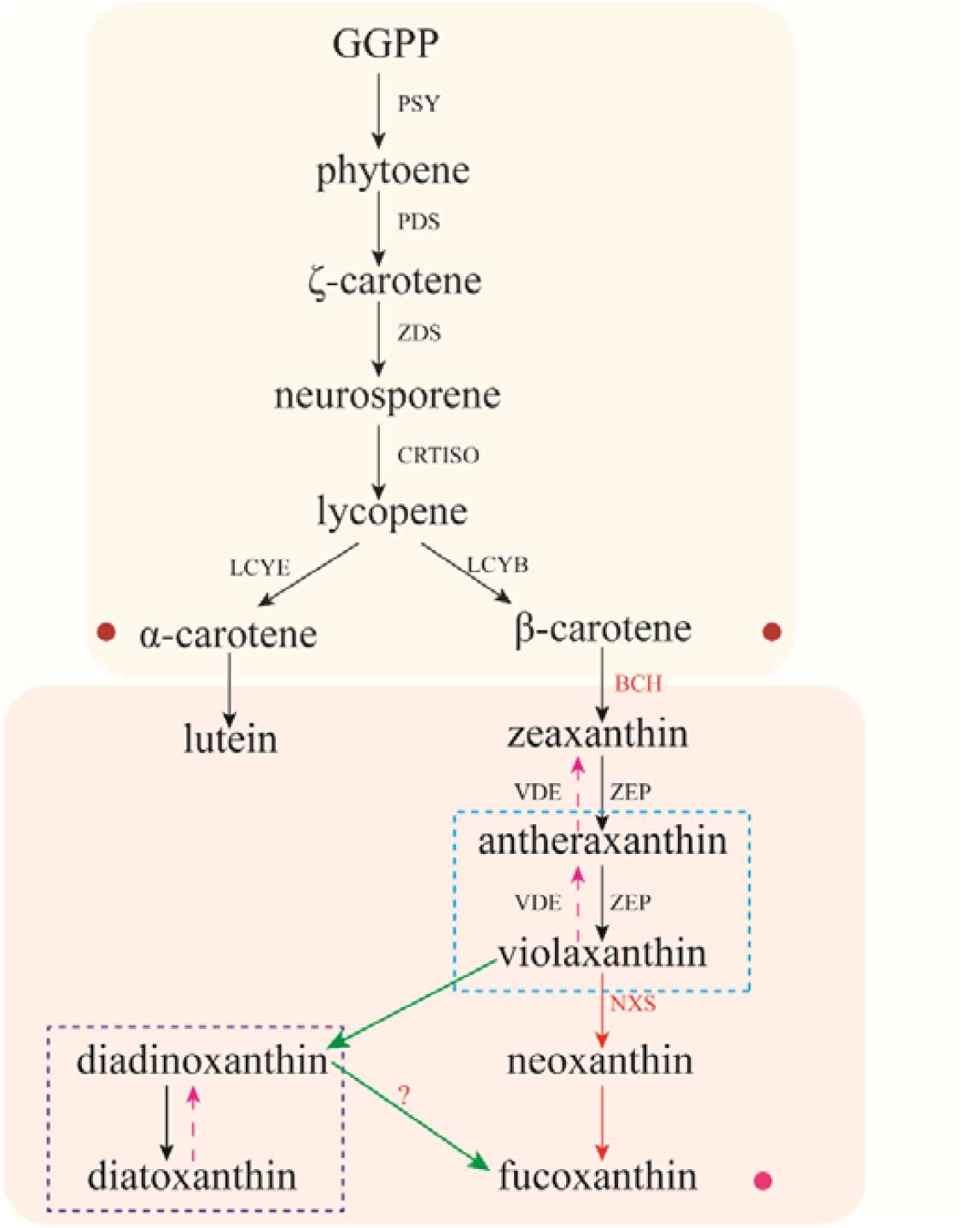
Proposed fucoxanthin biosynthesis steps in diatoms. The red arrows represent the hypothetical neoxanthin hypothesis pathways while the green arrows represent the diadinoxanthin synthesis pathway.

In fucoxanthin synthesis pathway, many key genes functioning from GGPP to β-carotene synthesis have been identified with their functions in carotenoid synthesis verified. However, most genes functioning in the steps from β-carotene to fucoxanthin remain unidentified although their homologs may exist in diatom genomes (Kuczynska et al., 2015). *Phaeodactylum tricornutum* is a model diatom with a well-characterized genome and engineering tools available. Researchers are devoted to identifying the possible genes involved in fucoxanthin biosynthesis. In the newest trial, the violaxanthin de-epoxidase-like (VDL) protein genes was cloned from *P. tricornutum*, which involved in the biosynthesis of neoxanthin from violaxanthin (Dautermann et al., 2020; Bai et al., 2022).

In this study, we evidenced the existence of neoxanthin in *P. tricornutum* with a trace amount of xanthophyll detection method and identified the neoxanthin synthase gene (*NXS*) from *P. tricornutum*. We also deduced the tentative pathway of fucoxanthin from violaxanthin in diatoms by CRISPR/Cas9-mediated knocking out the homologous gene of *P. tricornutum* and determining the abundance of its transcript.

## 2. Results

### 2.1 Growth of wild-type P. tricortutum and identification of its xanthophyll

Light intensity plays a crucial role in carotenoid metabolism (Gómez-Loredo et al., 2016; Nisar et al., 2015). To profile xanthophylls, we cultured the wide-type *P. tricornutum* in air-bubbled liquid medium (low CO_2_, *c*. 0.04% v/v) at 45 and 300 μmol m^-2^s^-1^ light intensities, respectively. Cell growth was characterized in terms of OD_678_ and biomass. Although the growth rate (OD_678_) was highly similar, the biomass under 45 μmol m^-2^s^-1^ was continuously higher than that under 300 μmol m^-2^s^-1^ during logarithmic growth phase (Fig. 2A). On day 10, the dry weight of the cells cultured under 45μmol m^-2^s^-1^ light intensity reached 2.1 g L^-1^, increased approximately 2-folds of that under 300 μmol m^-2^s^-1^ (Fig. 2B).

**Fig. 2.**
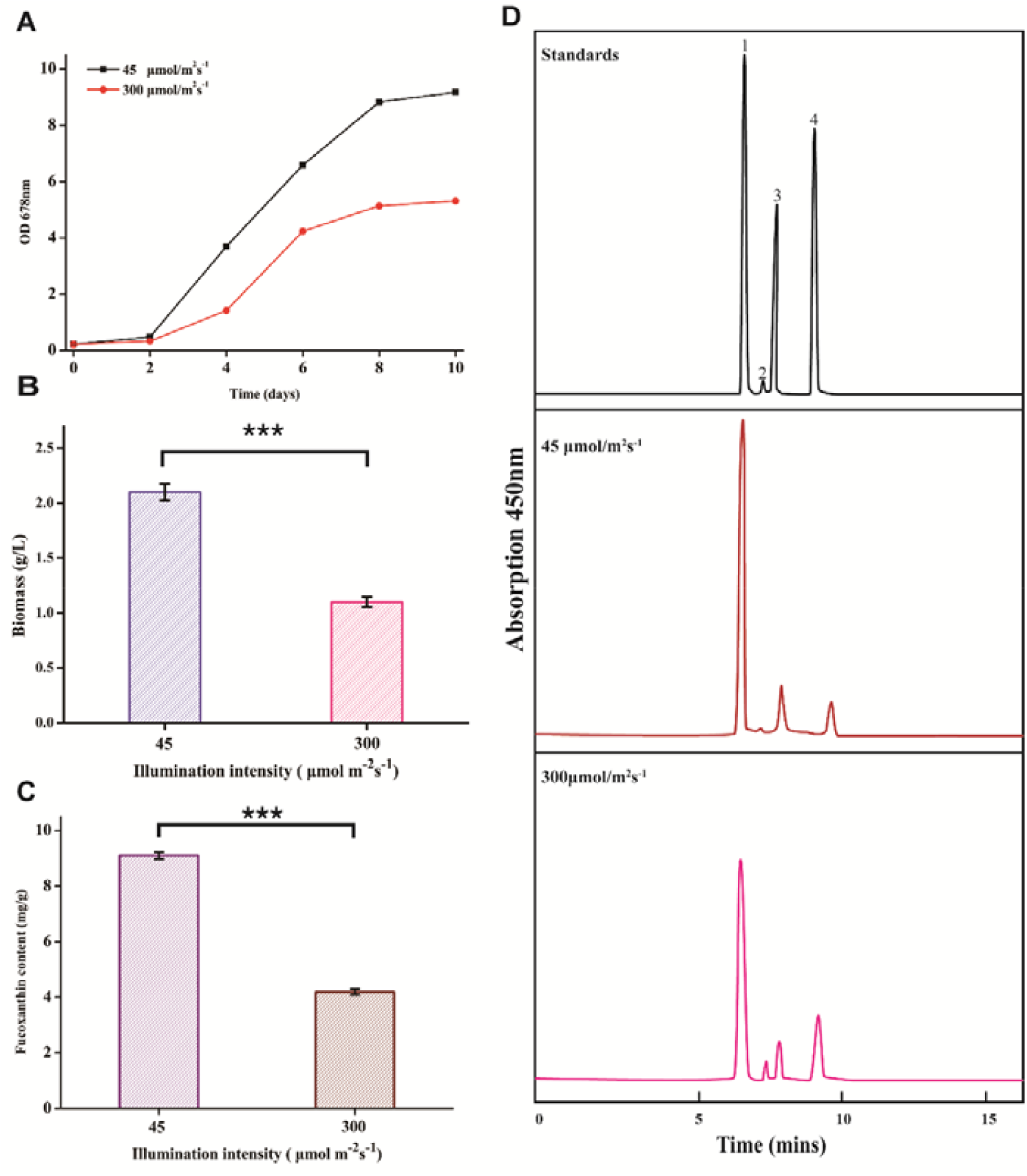
Growth and xanthophyll identification of wild-type *P. tricortutum*. Different light densities influenced the growth curve (A), biomass (B) and fucoxanthin content (C) of *P. tricornutum*. Carotenoids of *P. tricornutum* at different light densities were assayed at 450 nm (D). Peak 1, fucoxanthin; peak 2, neoxanthin; peak 3, zeaxanthin; peak 4, violaxanthin. Data are shown as means ± standard error (n=3). Asterisk triple indicates statistical significance (*p* < 0.05).

The carotenoids extracted from *P. tricornutum* were identified using high-performance liquid chromatography (HPLC) with a PDA detector. An example of xanthophyll distribution and profiles in concentrated fractions extracted from *P. tricornutum* was shown in Fig. 2C. The most abundant xanthophyll was fucoxanthin under both light intensities, being consistent with the previous report (Foo et al., 2017). Fucoxanthin reached 9.1 and 4.2 mg g^-1^ under 45 and 300 μmol m^-2^s^-1^, respectively, accounting for 0.9 % and 0.4% of the dry biomass (Fig. 2D). In addition, when separation time was extended, zeaxanthin and violaxanthin became more separatable and identifiable. However, their contents and changing trends were significantly different between two light intensities. Violaxanthin and zeaxanthin were 0.46 and 0.71 mg g^-1^ under 45 μmol m^-2^s^-1^. The content of zeaxanthin at high light intensity was 120% of that under low light intensity while the content of violaxanthin remained similar.

From the HPLC diagram, an obvious absorption peak at 7.4 min was observed in the extract of alga cultured under 300 μmol m^-2^s^-1^, which was extremely stronger than that observed in the extract of alga cultured at 45 μmol m^-2^s^-1^. Against the standards, we inferred that this peak corresponded to neoxanthin. The content of neoxanthin under high light intensity reached 25 mg kg^-1^, quite lower than that found in terrestrial plants (Zacarías-García et al., 2020). Such a scenario explained the failure of Bertrand in detecting microalgal neoxanthin (Bertrand, 2010). We successfully detected neoxanthin in the concentrated carotenoid extract of *P.tricornutum*. Adjusting the light intensity under which the alga was cultured changed the algal content of xanthophylls except violaxanthin. Low light intensity reduced the contents of zeaxanthin and neoxanthin, and enhanced the accumulation of fucoxanthin.

### 2.2 Isolation of neoxanthin synthase gene (NXS) of P. tricornutum

Neoxanthin synthase (NXS) catalyzing the convertion from violaxanthin to violaxanthin is common among terrestrial plants, green algae and red algae (Dambek et al., 2012). However, the gene coding NXS has not been successfully annotated in *P. tricornutum* genome (http://genome.jgi-psf.org/Phatr2). Diatoms originated from a secondary endosymbiotic event in which a nonphotosynthetic eukaryote probably engulfed a eukaryotic photosynthetic cell related to red algae with most algal genes transferred horizontally into the host nucleus (Keeling, 2004). We tried to isolate the homolog of algal *NXS* based on the similarity of sequences.

High-quality genomic DNA and cDNA obtained from *P. tricorntutum* cultured under 45 μmol m^-2^s^-1^ were used as templates for amplification of the diatom *NXS*. From the *NXS* of red alga *Cyanidioschyzon merolae* and the symbiotic dinoflagellate *Symbiodinium microadriaticum*, we designed different pairs of primers, and amplified diatom diatom *NXS* successfully (Fig. 3A). The PCR products were inserted into the expression vector pEASY-T1, and transferred into *E. coli* DH5α for sequencing.

**Fig. 3.**
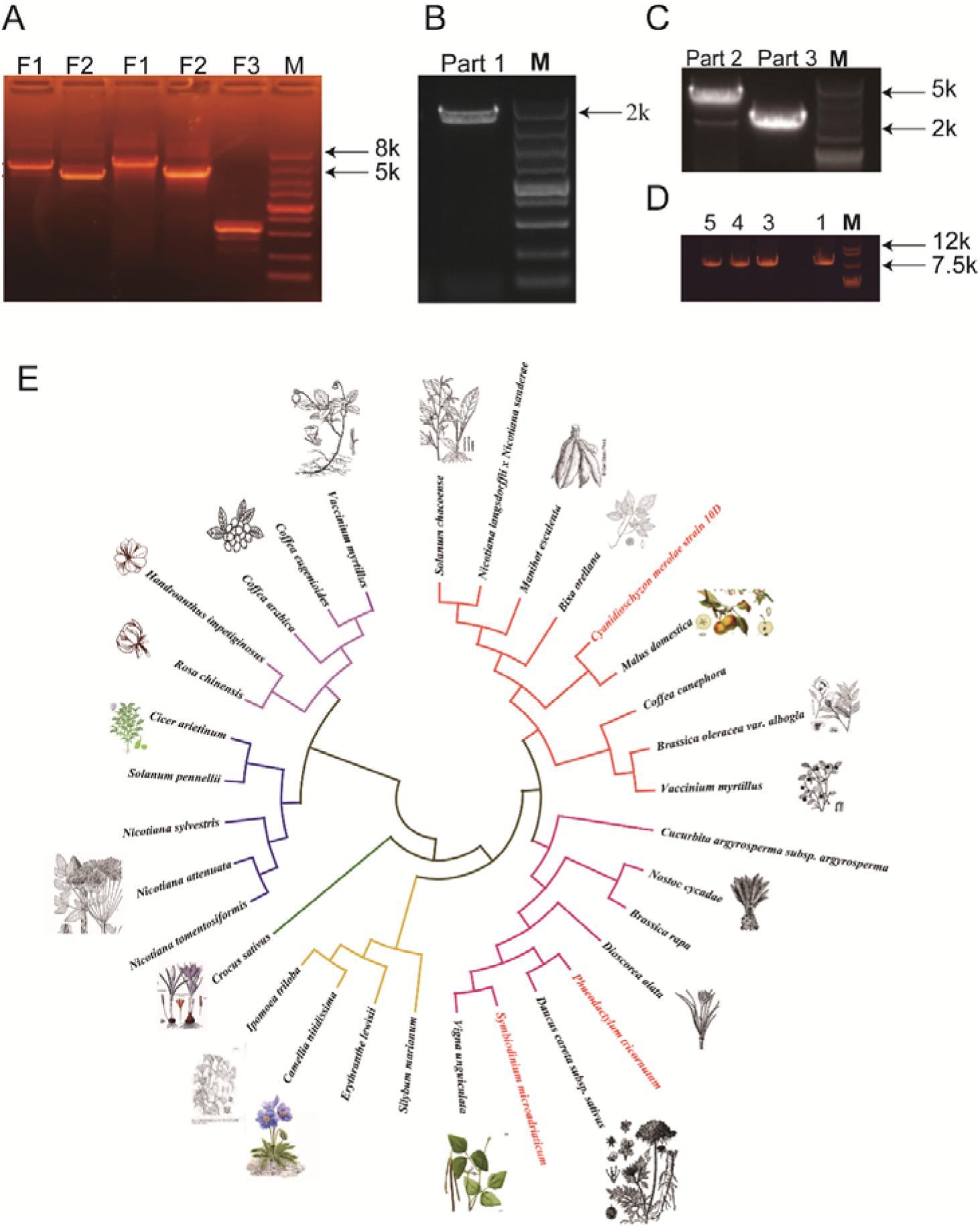
Isolation of *P. tricornutum NSX*. (A) the gene fragments amplified from genomic DNA and cDNA. (B) The fragment amplified with primer pair *NXS*1F/R. (C) The fragments amplified with primer pairs *NXS*2 F/R and *NXS*3 F/R. (D) the full length of *NXS* amplified with fusion PCR. (E) phylogenetic tree of the deduced diatom *NXS* and those retrieved form data banks.

We aligned the isolated sequence (XM_002177188.1) with those retrieved from data banks, and found that the similarity between the isolated and the downloaded was > 99%, indicagting that the isolated should be *NXS* of *P. tricortutum*. We designed the specific primers, *NXS*1,2,3*-*F and *NXS*1,2,3-R (Table S1) according to the CDS of XM_002177188.1, and amplified DNA and cDNA for 5’ and 3’ ends. We obtained three different PCR fragments (Figs. 3B and C), and merged them with fusion PCR (Fig. 3D). The gene is 8,265 bp in length while its cDNA is 8,193 bp in length. The cDNA sequence was deposited in GenBank with accession number MN525773.

The phylogenetic tree indicates that the cDNA of diatom *NXS* is highly homologous with *NXS* of *S. microadriaticum* and *Daucus carota* (Fig. 3E), *Daucus carota NXS* encodes neoxanthin synthase and functions in neoxanthin synthesis. The similarity between diatom *NXS* and *Daucus carota NXS* indicated that diatom *NXS* functions in diatom neoxanthin synthesis.

The abundance of *NXS* transcript under 45 and 300 μmol m^-2^s^-1^ light intensities was analyzed by real-time qPCR. We found that the expression of diatom *NXS* was up-regulated by 7-folds under high light intensity than that under low light intensity (Fig. 4C), which was paralleling with the accumulation of neoxanthin.

**Fig. 4.**
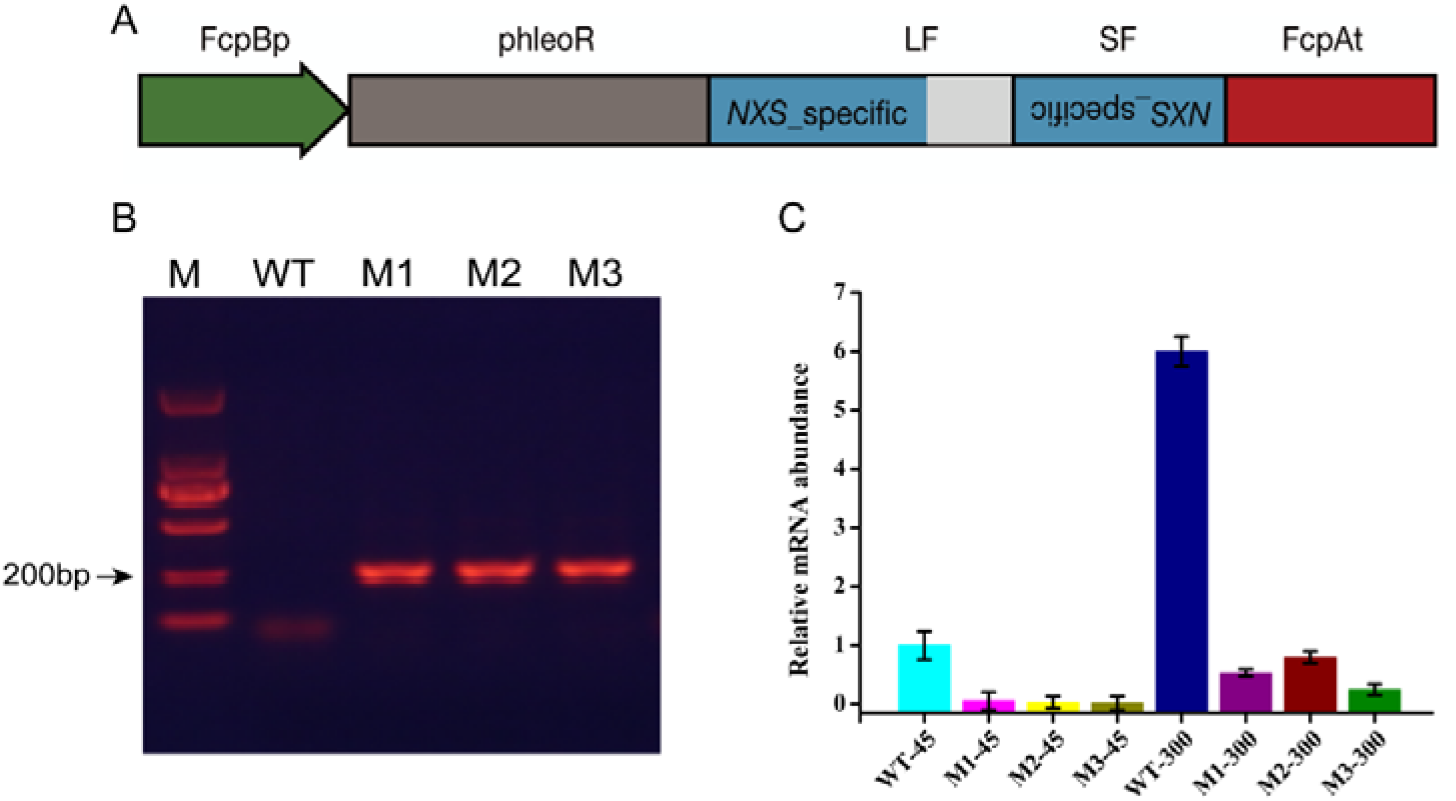
Generation and screening of the RNAi-based targeted gene knockdown mutant strains. (A) Expression cassette containing inverted repeats of the *NXS* gene. LF, long fragment; SF, short fragment; FcpBp: promoter of fucoxanthin chlorophyll protein gene from *P. tricornutum*; phleoR: Zeocin resistant gene; FcpAt: terminator of fucoxanthin chlorophyll protein gene from *P. tricornutum*. (B) Validation of the transgenic *P. tricornutum* lines by PCR. The lines were cultured in a liquid f/2 medium with zeocin for selection. (C) The transcript levels of *NXS* in strains under different light densities were determined by real-time qPCR, and calculated from Ct values using the 2^-ΔΔt^ method after all results were normalized against the β-actin housekeeping gene. Data are shown as means ± standard error (n=3).

### 2.3 Predicted structure and calculated characteristics of P. tricornutum NXS

Using software available online, we predicted the structure and calculated the characteristics of diatom NXS. The reveals by ProtParam included molecular weight, molecular formulas, instability coefficient and half-life (Fig. S1A). As predicted by SOPMA (Geourjon and Deléage, 1995), diatom NXS consisted of 50.37% of alpha-helix, 5.20% of beta-turn and 34.58% of random coil (Fig. S1B). Calculation with the Protscale tool (Xuan et al., 2013) showed that rich hydrophobic amino acids evenly distribute throughout the peptide chain (Fig. S1C). The TMHMM (Chen et al., 2003) predicted that diatom NXS has no transmembrane domain (Fig. S1D). The three-dimensional structure of diatom NXS is shown in Fig. S1E.

### 2.4 Knocking down P. tricornutum NXS expression

To decipher the function of *NXS* in fucoxanthin biosynthesis, RNA interference (RNAi) (Carthew and Sontheimer, 2009) was used in this study. We inverted a part of the gene to knock down the expression of diatom *NXS* as described early (Wei et al., 2017). A ∼200 bp short fragment (SF) and a ∼ 400 bp long fragment (LF) of the diatom *NXS* were amplified with primers listed in Table S1. These two fragments share the first 200 bp to ensure their transcripts will anneal into a partial double-stranded RNA which can be recognized by Dicer. The two fragments were independently amplified from *P. tricornutum* cDNA, oriented reversely and inserted into phir-PtNXS, generating a 5.35 kb recombinant plasmid (Fig. 4A), which contained FcpB promoter and selection marker *Sh ble*.

After biolistic transformation, zeocin-resistant colonies were picked and cultured. From the genomic DNA extracted from the cultured algal colonies, *Sh ble* was amplified to verify the successful transformation according to expected band (Fig. 4B).

Three *NXS* knockdown strains (M1, M2 and M3) were randomly selected and cultured under 45 and 300 μmol m^-2^s^-1^, respectively. The abundance of *NXS* transcripts each was determined through qPCR. As illustrated in Fig. 4C, the abundance of *NXS* transcript was 95-98 % and 86-96% lower than that of WT at two light intensities, respectively, indicating that the expression of diatom *NXS* was significantly down regulated.

### 2.5 Phenotype of P. tricornutum NXS knockdown strains

A comparison between *NXS* knockdown strains and WT at different light intensities (Fig. S2) showed that the growth rate of knockdown strains decreased slightly at both 45 and 300 μmol m^-2^s^-1^. Xanthophyll accumulated in strain M1 as was revealed with HPLC (Fig. 5A and B). At 300 μmol m^-2^s^-1^ light intensity, 0.007 mg g^-1^ of neoxanthin was detected in *NXS* knockdown strain, significantly lower than that in WT (0.025 mg g^-1^). Neoxanthin was not detectable in concentrated extract from *P. tricornutum* cultured under 45 μmol m^-2^s^-1^ light. Knocking down *NXS* decreased neoxanthin accumulation as we expected.

**Fig. 5.**
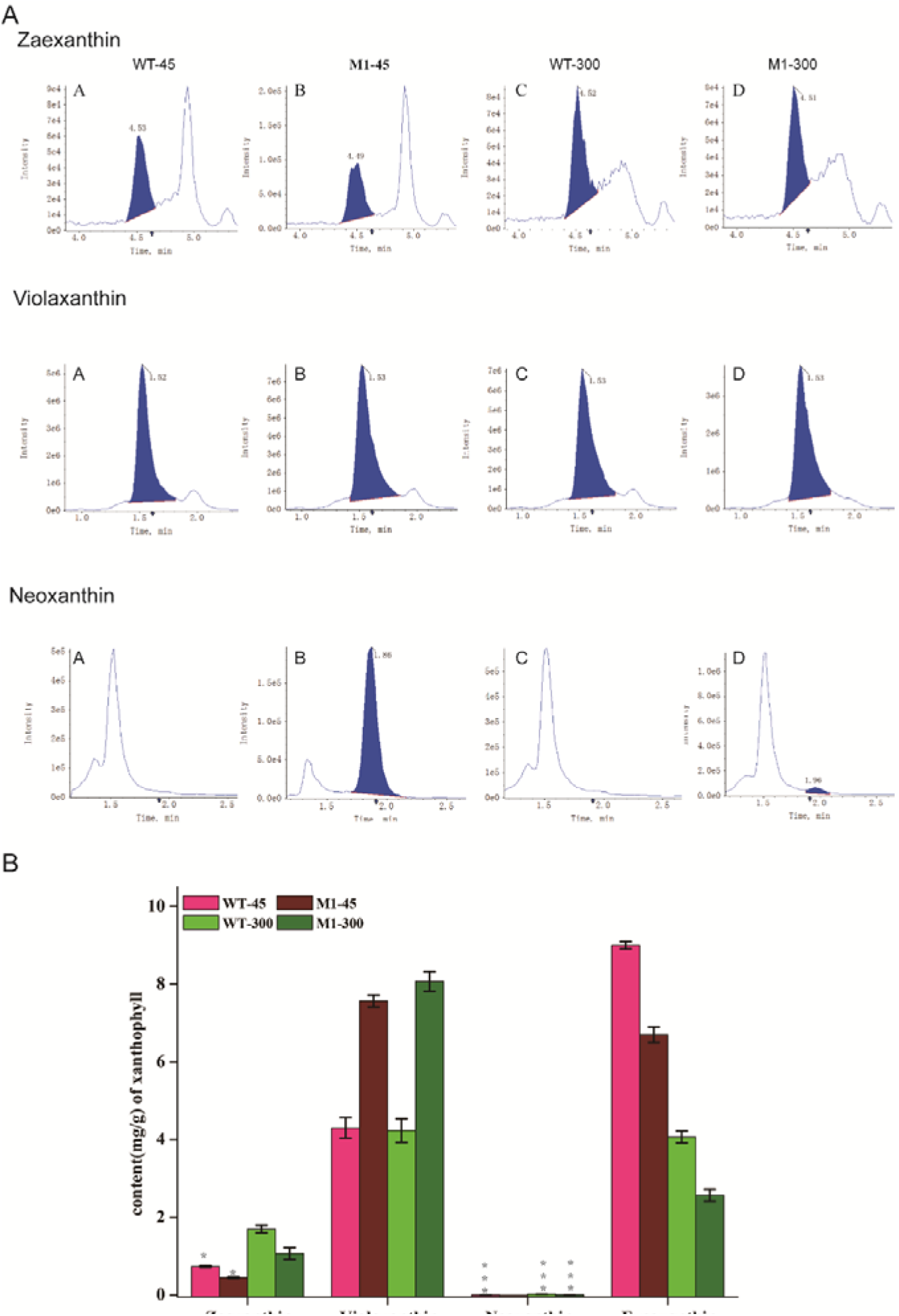
Comparsion of the pigment content in WT and *P. tricornutum NXS* knockdown strains. (**A)** Identification of carotenoids from *P. tricornutum* with HPLC. (**B)** The content of xanthophyll in knockdown strains and WT at different light intensities. Data are shown as means ± standard error (n=3).

Knocking down *NXS* expression also decreased the accumulation of zeaxanthin from 0.73 and 1.7 mg g^-1^ of WT to 0.44 and 1.06 mg g^-1^ of M1 under 45 and 300 μmol m^-2^s^-1^ irradiation, respectively. However, violaxanthin concentration in *NXS* knockdown strain reached 0.76 and 0.87 mg g^-1^ under 45 and 300 μmol m^-2^s^-1^ irradiation, respectively, 65% and 100% higher than that in WT.

Fucoxanthin remained as the major xanthophyll in M1. The accumulation of fucoxanthin in *NXS* knockdown strain was suppressed in comparison with WT at two light intensities. The concentration of fucoxanthin in *NXS* knockdown strain decreased to 6.7 and 2.6 mg g^-1^ at 45 and 300 μmol m^-2^s^-1^ light intensities, respectively, 24% and 38% lower than that of WT.

### 2.6 CRISPR/Cas9-mediated knockout of P. tricornutum NXS

Double-stranded DNA formed by the target oligonucleotides was inserted into pPtpuc3Cas9-sgRNA (Sharma et al., 2019) linearized at *Bsa* I (Karas et al., 2015), yielding pPtpuc3Cas9-sgRNA1 and pPtpuc3Cas9-sgRNA2. Phleomycin-resistant algal colonies were lysed to verify the insertion of recombinant plasmids by PCR. Ten pPtpuc3Cas9-sgRNA1 containing strainsand sixteen pPtpuc3Cas9-sgRNA2 strains (Fig. 6A) were picked, respectively. From all algal colonies, a band of about 250 bp in length was amplified. Two pPtpuc3Cas9-sgRNA1 strains and two pPtpuc3Cas9-sgRNA2 strains were selected for Western blotting assay. The expected hybridization band was found in transgenic strains but not founds in WT, indicating that the transformation was successful (Fig. 6B). PCR amplification and Sanger sequencing were conducted to identify the knockout sites. Nucleotide A was transformed into C in the Ko_1 strain (Fig. 6C) while T was deleted in the Ko_2 strain (Fig. 6D).

**Fig. 6.**
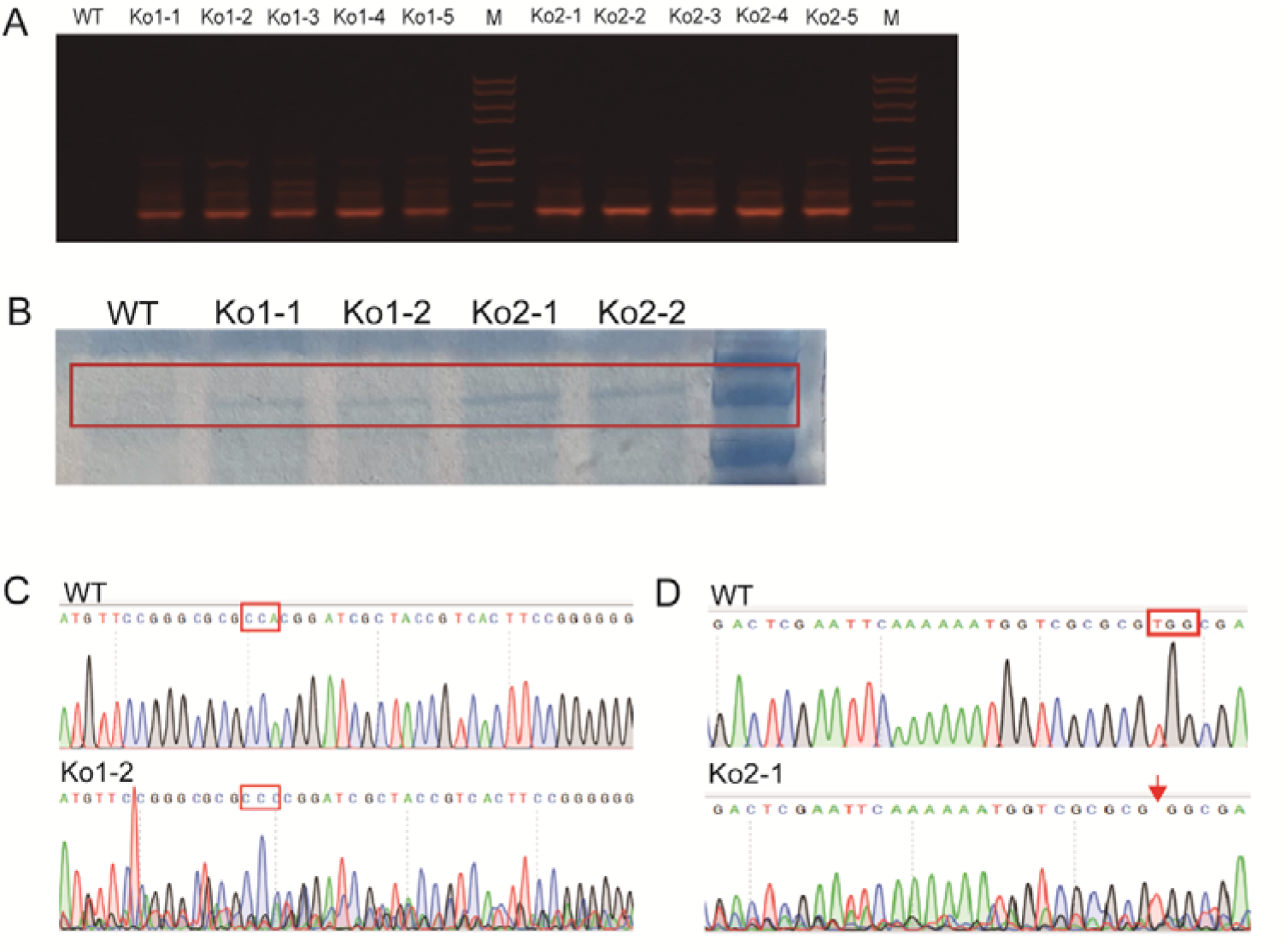
CRISPR/Cas9 mediated knocking out of *P. tricornutum NXS*. (A) Verification of transformants with PCR. As excepted, a 550 bp band was amplified among transgenic strains but not found in WT. (B) Varificartio of Cas9 thorough Western blotting. (C) and (D) Determination of CRISPR/Ca9 mediated knockout sites by sequencing.

### 2.7 Knocking out NXS affected xanthophylls profile and growth of P. tricornutum

The *NXS* knockout strains grew, but their growth rate was lowever than WT (Figs. 7A and B). In 10 days, the dry weight of the two strains was significantly lower than that of WT at both 45 and 300 μmol m^-2^s^-1^ (Figs. 7C and D). Knocking out *NXS* greatly impacted xanthophyll accumulation of *P. tricortutum* (Fig. 7E). NXS is the key enzyme for the synthesis of neoxanthin. When *NXS* is knocked out, no neoxanthin was detected in the concentrated extract of *P. tricornutum* cultured at both low and high light intensities. The content of zeaxanthin in Ko1-2 decreased to 0.33 and 0.41 mg g^-1^ at 45 and 300 μmol m^-2^s^-1^, respectively, 56% and 53% decreased in comparison with WT. Vioxanthin is the substrate of *NXS*, which significantly accumulated in Ko_1, reaching 0.88 and 0.95 mg g^-1^ at 45 and 300 μmol m^-2^s^-1^, respectively. However, the content of fucoxanthin significantly decreased to 4.1 and 2.2 mg g^-1^, respectively. These findings indicated that fucoxanthin synthesis pathway was blocked to a certain extent in *P. tricortutum* when the *NXS* was knocked out. Xanthophylls profile can be regulated by *NXS* in *P. tricornutum*. Decrease of *NXS* transcript abundance leads to the accumulation of zeaxanthin and reduction of violaxanthin, neoxanthin and fucoxanthin. We speculated that neoxanthin is indeed a key intermediate that is involved in the biosynthesis of fucoxanthin.

**Fig. 7.**
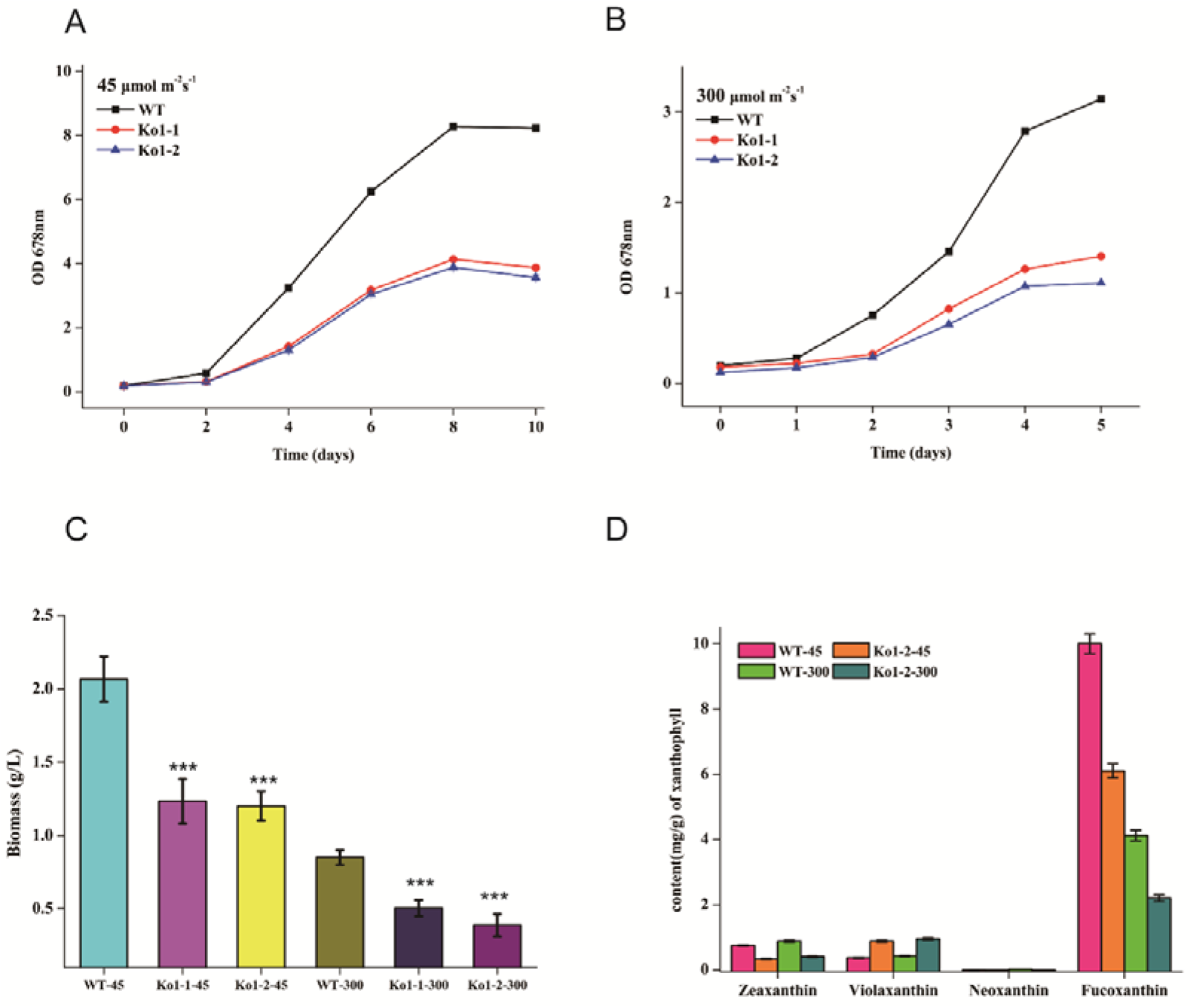
Growth performance and xanthophylls profile of WT and *P. tricornutum XNS* knockout strains. Growth curves **(A, B)** and biomass **(C)** changes of *NXS* knockout *P. tricornutum* strains at 45 μmol m^-2^s^-1^ and 300 μmol m^-2^s^-1^. **(D)** The content of xanthophylls in stranis at different light intensities. Data are shown as means ± standard error (n=3). Asterisk triple indicates statistical significance compaired with WT (*p* < 0.05).

## 3. Discussion

### 3.1 New evidence for neoxanthin and NXS existed in P. tricornutum

The fucoxanthin biosynthetic pathway in diatom has drawn great attention for its important physiologic functions. Violaxanthin, one of the intermediates, was previously regarded as a precursor of fucoxanthin in *P. tricornutum* (Lohr and Wilhelm, 2001). The biosynthetic pathway from GGPP to violaxanthin has been elucidated based on the genome information and molecular tools available (Eilers et al., 2016). However, the conversion of violaxanthin to fucoxanthin remains unclear, and the role of neoxanthin is key to probing the reactions from violaxanthin to fucoxanthin. Neoxanthin is common in terrestrial plants and red algae (Kakinuma et al., 2001). However, the neoxanthin biosynthetic pathway in diatom is debatable cause neoxanthin can not be detected in most cases. In our study, neoxanthin was not detected in standard carotenoid extracts from *P. tricornutum* at first, which was consistent with the previous study (Kim et al., 2012). Nevertheless, after further enriching and concentration, the absorption peak of neoxanthin in the carotenoid extract was detected successfully (Fig. 2C). Although the neoxanthin content is at ppm level, the presence of neoxanthin in diatom can be confirmed.

In addition, quantitative analysis revealed that violaxanthin kept a stable content under different light intensities while the content of neoxanthin and fucoxanthin changed significantly under different light intensities (Havaux and Niyogi, 1999). The violaxanthin cycle and diadinoxanthin cycle constitute two short-term photoprotective mechanisms in diatoms, and neoxanthin and fucoxanthin participate in these two cycles respectively (Goss et al., 2007; Lepetit et al., 2010; Havaux and Niyogi, 1999). When light intensity increases, neoxanthin tends to be converted for self-protection, leading to a decrease in fucoxanthin synthesis (Wilhelm et al., 2006). According to the relationship between neoxanthin and fucoxanthin concentration variation under different light intensities, we infer that neoxanthin acts as an intermediate in fucoxanthin biosynthesis in *P. tricornutum*.

To confirm that neoxanthin was synthesized from violaxanthin, we aligned the multi-sequences of neoxanthin synthetase (NXS) deduced from other algae and identified their conserved sequences for primers designing. We cloned the full-length *NXS* gene (accession number MN525773) in *P. tricornutum* successfully. qPCR analysis shows that the expression of *NXS* can be regulated by light intensity, and is consistent with the accumulation of neoxanthin.

### 3.2 NXS regulates the accumulations of xanthophylls

Although many genes involved in carotenoid biosynthesis have been tentatively identified and summarized, there are still few reports about the putative genes involved in the conversion of violaxanthin to neoxanthin. Shen et al. identified a gene named *PtABA4* that has a potential function relevant to violaxanthin conversion (Shen et al., 2022). However, the change in neoxanthin accumulation was not observed with the expression of *PtABA4*. Dautermann et al. cloned violaxanthin de-epoxidase-like genes (*VDL*) from *P. tricornutum* and characterized them as the key enzymes involved in the conversion of violaxanthin to neoxanthin. However, the genes are functionally verified in tobacco leaves, not in *P. tricornutum* (Dautermann et al., 2020).

In our study, *P. tricornutum NXS* knockdown and knockout strains were obtained to further probe the role of *NXS* in the biosynthetic pathway of fucoxanthin. The accumulations of neoxanthin, zeaxanthin, and fucoxanthin were decreased in *NXS* knockdown strains which were positively correlated with the expression levels of *NXS*. The suppression of the *NXS* gene results in the rise of violaxanthin and the decline of neoxanthin in *P. tricornutum*, and no neoxanthin was detected in the *NXS* knock-out mutant. These results confirmed that *NXS* (accession number MN525773) is involved and may catalyze the conversion of neoxanthin from violaxanthin. *NXS* is totally different from *VDL* and *PtABA4* in the former literature. Meanwhile, the accumulation of fucoxanthin shows varying degrees of reduction along with the decline of *NXS* expression and neoxanthin content, revealing that fucoxanthin is a downstream product of neoxanthin.

### 3.3 The deduced biosynthetic pathway of fucoxanthin from violaxanthin in diatom

All modifications to the conversion of violaxanthin from β-carotene are symmetrical, i.e. both ends of the carotenoid molecule are modified in the same ways. Asymmetrical modifications start with the formation of allenic double bonds at only one side of the carotenoid molecule forming neoxanthin. This reaction proceeds via an opening of one 5, 6-epoxy ring and the rearrangement to allenic double bonds at position 6 of the β-ionone ring. From the molecular structure analysis of neoxanthin, diadinoxanthin, and fucoxanthin, further conversion from neoxanthin to diadinoxanthin can be formed in a single reaction by the formation of an acetylenic bond from the allenic double bonds by elimination of the 5-HO group as water (Coesel et al., 2008). Meaningfully, our research offers further actual data evidence in a sequential reaction of violaxanthin conversion to neoxanthin, and then diadinoxanthin.

As follows, we find that the formation of fucoxanthin could not be possibly conversed in one reaction from either neoxanthin or diadinoxanthin, due to both ends of the molecular structure of fucoxanthin being changed. Therefore, there should be multiple enzymes or complex enzymes involved in the conversion. Otherwise, there might be other intermediate xanthophylls in the process of fucoxanthin from either neoxanthin or diadinoxanthin. Meanwhile, in our study, when the *NXS* gene was knockout, the synthesis of fucoxanthin was indeed decreased but not blocked, which inferred that neoxanthin was not the only branch point of fucoxanthin and diadinoxanthin. There might be another synthetic pathway from violaxanthin to fucoxanthin in *P. tricornutum*, it could be diadinoxanthin or something else.

The biosynthetic pathway from violaxanthin to fucoxanthin is very complex, with a diverse range of intermediate xanthophylls involved in the process. Thus it is meaningful to detecte more kinds of xanthophylls and identificate genes encoding ketolase or acetylase in further study.

## 4. Methods

### 4.1 Microalgae strains and growth conditions

Cells were inoculated in a 100 mL Erlenmeyer flask containing 30 mL of sterile f/2 medium (Guillard., 1975) at 23 ± 1 °C under an illumination intensity of 30 μmol m^-2^s^-1^. The flasks were shaken manually twice a day. Pellets were harvested via centrifugation and washed with distilled water twice, and then were grown in a 500 mL column with 400 mL f/2 medium. Compressed air with 0.04% CO_2_ was pumped into the bottom of the column to supply the carbon source and mix the culture. Cultures were cultivated under 45 and 300 μmol m^-2^s^-1^ continuous white light, respectively.

The optical densities (OD) of algae cells under different light intensities were calculated with a U-3310 spectrophotometer (Hitachi, Toyko, Japan). Moreover, for dry weight measurement, aliquots (5 mL) of the cell culture were filtered through Whatman GF/C paper, washed three times with distilled water, and dried at 105 °C for 4 h. All the trials were repeated at least 3 times.

### 4.2 Pigments extraction and analysis

For total carotenoid pigments extraction, cells collected by centrifugation were immediately frozen in liquid nitrogen and then freeze-dried in a freeze-dryer. Lyophilized algae samples (10 mg) were added to 1 mL using cold methanol/acetone (1:1, v/v) solution and homogenized using a homogenizer. Then, the pigments were extracted on ice (250 rpm) in the dark. An hour later, the samples were centrifuged at 4,000 g for 10 min to collect the supernatant. Such an extraction process was repeated 3 times, and the supernatants were filtered by a 0.22 μm PTFE filter before conducting further analysis. Only for enrichment, carotenoid extracts were fractionated and concentrated by thin-layer chromatography (TLC) on activated silica plates developed with cold methanol/acetone (1:1, v/v) solution, and the supernatants were filtered by a 0.22 μm PTFE filter.

Pigments were detected through Waters 2695 HPLC systems (Waters, Milford, MA, USA) with a UV or PDA detector and a C18 reverse-phase bar (5 mm particle size, 250 × 4.6 mm). The mobile phase used was (A) 85% methanol and (B) 100% ethyl acetate, with a flow rate of 0.8 ml min^-1^. Pigment absorption spectra were collected from 210 to 700nm. Peaks for carotenoids were identified according to their absorption at 449nm (Guo et al., 2016).

### 4.3 Gene isolation and bioinformatic analysis of NXS

DNA and cDNA of *P. tricornutum* were used as templates for PCR with TransStart FastPfu (TransGen Biotech, Beijing, China) polymerase. Different pairs of primers were designed based on the *NXS* gene of red algae *Cyanidioschyzon merolae* strain 10D (F1) and the symbiotic dinoflagellate *Symbiodinium microadriaticum* (F2, F3) (Table S1). Aligned the sequence of each PCR product in NCBI and found that all PCR products had over 99% similarity with the sequence (accession number XM_002177188.1) which was an unknown gene from *P. tricornutum*.

In order to obtain the full sequence of *NXS*, three pairs of primers used are listed in Supplementary Table 1 to clone *NXS* part 1, part 2, and part 3. The fusion PCR was used to ligate these parts and obtain full-length *NXS*. The curated alignment was then used to construct a phylogenetic tree with the neighbor-joining method in MEGA11. Bootstrap value, the percentage of replicate trees in which the associated taxa clustered together was tested for 1,000 replicates.

The physical property of the candidate *NXS* sequence from *P. tricornutum* was analyzed by an online software ProtParam (Garg et al., 2016), the Target-P program for the prediction of possible plastid localization, and the ChloroP1.1 server (Gasteiger, 2005) for the identification of a chloroplast transit peptide. For transmembrane domain analysis, the TMHMM server (Krogh et al., 2001) was used. The domain structure of the predicted protein was analyzed by PyMol (Delano, 2002).

### 4.4 Construction of vectors

For the *NXS* RNAi vector, a 200-bp short fragment (corresponding to the nucleotide sequence from 71 to 270 bp) and a 350-bp long fragment (corresponding to the *NXS* sequence from 71 to 420 bp) were amplified from the *P. tricornutum* cDNA, respectively, with the primers nxs400-Insert1-Primer F and nxs400-Insert1-Primer R, and nxs200-Insert1-Primer F and nxs200-Insert1-Primer R (Supplementary Table 1). These two fragments had the first 200 bp in common, ligated in sense and antisense orientations to the linearized Pt-NXS vector digested with *Eco*R I to create phir-PtNXS plasmid by using the MultiS One Step Cloning Kit (Vazyme, Nanjing, China) according to the manufacturer’s instructions.

For the *NXS* knockout vector, we used a custom-made or publicly available CRISPR/Cas9 target finding tool (e.g., PhytoCRISP-Ex) to identify Cas9 target sites (N20-NGG to) with low or no homology to other genomic loci. Order 24 nt long oligos with 20 complementary nt and 5’ TCGA and AAAC overhangs for the creation of the adapter for targeting the gene of interest (Supplementary Table 1). Exploit the two *Bsa* I restriction sites that are placed immediately 5’-to the sgRNA in the vector to ligate the adapter with 5’-TCGA and 5’-AAAC overhangs into the pPtpucCas9-sgRNA plasmid. Each plasmid sequence was verified by sequencing before transformation into *P. tricornutum*.

### 4.5 Transformation of P. tricornutum

RNAi vector was transformed into WT *P. tricornutum* by biolistic transformation according to (Apt et al., 1996) with minor modifications. The bombardment was performed (a distance of 6 cm using 1350 psi rupture disks) with a PDS-1000/He Biolistic Particle Delivery system (Bio-Rad, USA). The antibiotic concentration was determined after a sensitivity test, optimal screening conditions were 70 μg ml^-1^ of zeocin.

The pPtPuc3m diaCas9_sgRNA plasmid was delivered to *P. tricornutum* cells via conjugation from *E coli* DH10β cells as described (Sharma et al., 2019). *E.coli* DH10β cells with mobilization helper plasmid pRL443, containing all genes necessary for the conjugative transfer of oriT-containing plasmids. *E. coli* pRL443 competent cells were transformed with pPtPuc3m diaCas9_sgRNA plasmid and used for conjugative delivery of the CRISPR-Cas9 plasmid to the cells. Once the plasmid is transformed into *P. tricornutum* will replicate as an episome and remain stable if propagated in zeocin-containing growth media.

### 4.6 Quantitative qRT-PCR

The relative abundance of *P. tricornutum NXS* gene was examined by qRT-PCR on an IQ5 Real-Time PCR Detection System. In each experiment, a specific concentration of a PCR fragment was amplified in 20 μL of reaction solution containing 1 × SYBR Green PCR Master Mix and corresponding primers for *NXS*. Data were captured as amplification plots. Transcription levels of the target gene were calculated from the threshold cycle by interpolation from the standard curve (Livak and Schmittgen, 2001). The actin gene was also used as the internal standard for *NXS* mRNA analysis in *P. tricornutum*. The complete experiments (RNA isolation, cDNA synthesis followed with qRT-PCR) were repeated twice independently, and data are the average of at least three replicates.

### 4.7 SDS-PAGE and Western-Blot Analysis

For protein analysis, cells grown under 45 and 300 μmol m^-2^s^-1^ light intensities were harvested, and then resuspended in lysis buffer (20□mM HEPES-KOH pH 7.5, 5□mM MgCl_2_, 5□mM β-mercaptoethanol, and 1□mM PMSF) and disrupted by sonication. Total protein samples were loaded and separated on an SDS-PAGE gel. The SDS-PAGE gels were stained with 0.1% (w/v) Coomassie Brilliant Blue R for visualization or blotted onto an ATTO P PVDF membrane via a semi-dry transfer system. Membranes were probed with specific polyclonal antibodies raised against the Cas9.

### 4.8 Statistical Analysis

All cultivations were performed at least in duplicate. All determinative data were collected from triplicate samples and the final values were expressed as mean value ± standard deviation. Statistical significance was evaluated by ANOVA and t-test using SPSS programs (Version 19.0, IBM SPSS, Chicago IL, USA) at a level of *P* <0.05.

## Author contributions

S.J.Kang and H.Wang initiated the program, coordinated the project, and wrote the manuscript. S.J.Kang, Z.Y.Zhang, and J.C.Han prepared and analyzed the samples. G.P. Yang and T.Z. Liu gave instructions during the research and revised the manuscript.

## Acknowledgments

This work was financially supported by the National Key Research and Development Program of China (Grant No.2018YFA0902500) and Shangdong Taishan Scholars Program (No. tsqn202103144).

## Supplementary data

Reference genome data are deposited in GenBank with the number MN525773.

Table. S1. All different pairs of primers for this experiment.

Fig.1. Physical property and molecular structure of NXS in *P. tricornutum*

Fig.2. Phenotype of the *P. tricornutum* NXS knockdown mutants.

